# A hyper-immunogenic and slow-growing fungal strain induces a murine granulomatous response to cryptococcal infection

**DOI:** 10.1101/2021.10.26.466037

**Authors:** Calla L. Telzrow, Shannon Esher Righi, Natalia Castro-Lopez, Althea Campuzano, Jacob T. Brooks, John M. Carney, Floyd L. Wormley, J. Andrew Alspaugh

## Abstract

Many successful pathogens cause latent infections, remaining dormant within the host for years but retaining the ability to reactivate to cause symptomatic disease. The human opportunistic pathogen *Cryptococcus neoformans* is a ubiquitous yeast that establishes latent pulmonary infections in immunocompetent individuals upon fungal inhalation from the environment. These latent infections are frequently characterized by granulomas, or foci of chronic inflammation, that contain dormant cryptococcal cells. Immunosuppression causes these granulomas to break down and release viable fungal cells that proliferate, disseminate, and eventually cause lethal cryptococcosis. This course of *C. neoformans* dormancy and reactivation is understudied due to limited models, as chronic pulmonary granulomas do not typically form in most mouse models of cryptococcal infection. Here, we report that a previously characterized *Cryptococcus*-specific gene which is required for host-induced cell wall remodeling, *MAR1,* inhibits murine granuloma formation. Specifically, the *mar1*Δ loss-of-function mutant strain induces mature pulmonary granulomas at sites of infection dormancy in mice. Our data suggest that the combination of reduced fungal burden and increased immunogenicity of the *mar1*Δ mutant strain stimulates a host immune response that contains viable fungi within granulomas. Furthermore, we find that the *mar1*Δ mutant strain has slow growth and hypoxia resistance phenotypes, which may enable fungal persistence within pulmonary granulomas. Together with the conventional primary murine infection model, latent murine infection models will advance our understanding of cryptococcal disease progression and define fungal features important for persistence in the human host.

## INTRODUCTION

Granulomas are complex foci of chronic inflammation that form in response to many stimuli, including microbial infections. A hallmark of indolent infections such as tuberculosis disease, granulomas are often characterized by epithelioid macrophages, multinucleated giant cells, and dormant and/or slowly proliferating microorganisms (1–4). The traditional understanding of the granuloma considered it to be a host-directed defense response that restricts microbial access to nutrients and oxygen, resulting in an immune microenvironment that limits microbial proliferation and prevents dissemination (3, 4). However, more recent work has demonstrated that granulomas are a dynamic component of the complex host-microbial “arms race”. In addition to serving as a host-directed protection mechanism, microorganisms can exploit the granuloma as a micro-niche for long-term survival in the host, where they remain shielded from immune detection until microbial reactivation (3–5). Although most work on granulomas has been conducted in the context of mycobacterial infections, many other infectious microorganisms induce granuloma formation in the human lung (6).

The fungal pathogen *Cryptococcus neoformans* is a significant cause of pneumonia and fatal meningoencephalitis in immunocompromised populations around the world, resulting in more than 180,000 deaths annually (7). Primary infection occurs upon inhalation of environmental *C. neoformans* cells and/or spores, often early in life (4, 8). Immunocompetent hosts typically control the primary infection, with fungi remaining dormant but viable within lung-associated granulomas (9). As a result, immunocompetent hosts do not manifest infection-related symptoms with disease during this stage of latency (10). However, this latent infection can reactivate when a previously exposed individual becomes immunocompromised, especially in the setting of CD4+ T cell functional deficiency due to HIV infection, organ transplantation, and immunosenescence (2, 4, 7, 11). Breakdown of the cryptococcal granuloma structure results in microbial proliferation and systemic dissemination, including to the central nervous system.

The reactivation of fungal cells from granulomas is an understudied facet of cryptococcal disease, largely due to limited reactivation models. Although the mouse is the most well-characterized and commonly used animal model to study *Cryptococcus*-host interactions, many murine models do not form sustained granulomas in response to clinically relevant isolates of *C. neoformans* (12). As a result, most murine experiments focus on primary cryptococcal infection and subsequent systemic dissemination. To explore cryptococcal latency and reactivation, investigators have adopted models of cryptococcosis in rabbits (13) and rats (14, 15) or employed less virulent *C. neoformans* strains in mice (12, 16, 17). Recently, a novel latent model was reported in which pulmonary granulomas form in mice in response to infection with the *gcs1*Δ mutant cryptococcal strain lacking the glucosylceramide synthase (18–21). Mimicking the typical course of human disease, *gcs1*Δ cells induce well-formed granulomas in the lungs which contain dormant *gcs1*Δ cells that become reactivated from granulomas and disseminate upon immunosuppression (22).

We recently reported the identification and characterization of the *C. neoformans MAR1* gene that is required for cell surface remodeling in response to the host environment (23). The *mar1*Δ loss-of-function mutant strain displays altered cell surface features when exposed to physiological conditions, including decreased cell wall glucans and mannans, increased exposure of cell wall chitin, and impaired polysaccharide capsule attachment (23). The cell surface alterations of the *mar1*Δ mutant strain result in enhanced macrophage activation *in vitro* and hypovirulence in a murine inhalation model of cryptococcosis (23). We report here that this hyper-immunogenic *mar1*Δ mutant strain induces pulmonary granulomas in mice, resulting in a chronic and indolent infection. Furthermore, we describe both fungal and host factors that contribute to this granuloma response. From the fungal perspective, the combination of reduced fungal burden and hyper-immunogenicity of the *mar1*Δ mutant strain stimulates a host immune response that contains *mar1*Δ mutant cells within well-circumscribed granulomas. From the host perspective, we find that host GM-CSF signaling, a known contributor to granuloma formation (17, 24–26), is required for the formation of these granulomas. Finally, *in vitro* studies demonstrate that the *mar1*Δ mutant strain has cell cycle defects that may contribute to a slow growth phenotype and hypoxia resistance, two features which likely enable cryptococcal persistence within pulmonary granulomas. Because *MAR1* is a *Cryptococcus*-specific gene, this model represents a unique addition to the limited tools available to study the reactivation model of cryptococcal disease.

## MATERIALS & METHODS

### Strains, media, and growth conditions

All strains used in this study were generated in the *C. neoformans* var. *grubii* H99 (*MAT*α) (13) background and are included in Table 1. Strains were maintained on yeast extract-peptone-dextrose (YPD) medium (1% yeast extract, 2% peptone, 2% dextrose, and 2% agar for solid medium). Unless otherwise indicated, strains were incubated at 30°C.

**Table 1.** Fungal strains used in this study.

| Strain | Genotype | Source |
| --- | --- | --- |
| H99 | *MATα* | (13) |
| MAK1 | *MATα mar1Δ::NAT* | (23) |
| MAK11 | *MATα mar1Δ::NAT + MAR1-NEO* | (23) |
| cap59 | *MATα cap59Δ::NEO* | (63) |
| HEB6 | *MATα sre1Δ::NEO* | (31) |

### Histology analyses

The murine inhalation model of cryptococcosis was exclusively used in this study (27). For initial histological examination, C57BL/6 female mice were acquired from Charles River Laboratories. Mice were anesthetized with 2% isoflurane utilizing a rodent anesthesia device (Eagle Eye Anesthesia, Jacksonville, FL) and were infected via the intranasal route with 1 × 10^4^ CFU of either the wild-type (WT) (H99) or the *mar1*Δ mutant (MAK1) strain. Mice were sacrificed at predetermined endpoints (3, 7, 14, and 40 DPI) by CO_2_ inhalation followed by an approved secondary method of euthanasia. Lungs were perfused with and stored in 10% neutral buffered formalin. Lungs were subsequently paraffin-embedded, sectioned, mounted, and stained with hematoxylin and eosin by the Duke University School of Medicine Research Immunohistochemistry Shared Resource.

To determine the role of GM-CSF signaling in granuloma formation in this model, lungs from male and female Csf2rb^-/-^ mice (The Jackson Laboratory # 005940) were prepared as described above, with a few alterations. Mice were sacrificed at the predetermined endpoints of 3, 7, and 14 DPI by CO_2_ inhalation followed by an approved secondary method of euthanasia and lungs were perfused with PBS. The right lung was stored in 10% neutral buffered formalin for future histopathology preparation, while the left lung was used for fungal burden quantification analyses, as described below.

### Fungal burden quantification

Mice were infected as described above. Mice were euthanized at predetermined endpoints by CO_2_ inhalation followed by cervical dislocation, and lung tissues and/or brain tissues were excised. The left lobe of the lung and/or the brain was removed and homogenized in 1 mL of sterile PBS as previously described (28) followed by culture of 10-fold dilutions of each homogenate on YPD agar medium supplemented with chloramphenicol. Colony-forming units (CFU) were enumerated following incubation at 30°C for 48 hours. Statistical significance was determined using Student’s *t* test (GraphPad Software, San Diego, CA).

### Pulmonary cytokine analyses

C57BL/6 female mice acquired from Charles River Laboratories were infected and sacrificed as described above. Cytokine levels within the lung homogenates of infected mice were analyzed using the Bio-Plex protein array system (Luminex-based technology, Bio-Rad Laboratories, Hercules, CA). Briefly, lung tissues were excised and homogenized in 1 mL ice-cold sterile PBS. An aliquot (50 µl) was taken to quantify the pulmonary fungal burden, and an anti-protease buffer solution (1 mL) containing PBS, protease inhibitors, and 0.05% Triton X-100 was added to the homogenate. Samples were then clarified by centrifugation (3,500 rpm) for 10 minutes. Supernatants from pulmonary homogenates were assayed for the presence of IL-1α, IL-1β, IL-2, IL-3, IL-4, IL-5, IL-6, IL-9, IL-10, IL-12 (p40), IL-12 (p70), IL-13, IL-17, KC (CXCL1), MCP-1 (CCL2), MIP-1α (CCL3), MIP-1β (CCL4), RANTES (CCL5), Eotaxin (CCL11), IFN-γ, tumor necrosis factor (TNF)-α, granulocyte macrophage-colony stimulating factor (GM-CSF), and granulocyte-colony stimulating factor (G-CSF) according to the manufacturer’s instructions. Statistical significance between strains at each timepoint was determined using Student’s *t* test (GraphPad Software, San Diego, CA).

### Pulmonary leukocyte isolation

C57BL/6 female mice acquired from Charles River Laboratories were infected and sacrificed as described above. Lungs of infected mice were excised on 1, 3, 7, 14, and 21 DPI as previously described (28). Lungs were then digested enzymatically at 37°C for 30 minutes in 10 mL digestion buffer (RPMI 1640 and 1 mg/mL collagenase type IV [Sigma-Aldrich, St. Louis, MO]) with intermittent (every 10 minutes) stomacher homogenizations. The digested tissues were then successively filtered through sterile 70- and 40-μm nylon filters (BD Biosciences, San Diego, CA) to enrich for leukocytes, and the cells were then washed three times with sterile Hank’s Balanced Salt Solution (HBSS). Erythrocytes were lysed by incubation in NH_4_Cl buffer (0.859% NH_4_Cl, 0.1% KHCO_3_, 0.0372% Na_2_EDTA [pH 7.4]; Sigma-Aldrich) for 3 minutes on ice followed by a 2-fold excess of sterile PBS.

### Flow cytometry analyses

Pulmonary leukocytes were isolated from infected mice as described above. Standard methodology was employed for the direct immunofluorescence of pulmonary leukocytes (28, 29). Briefly, in 96-well U-bottom plates, 100 µl containing 1 × 10^6^ cells in PBS were incubated with yellow Zombie viability dye (1:1000 dilution, Cat. No 423104, Biolegend, San Diego, CA) for 15 minutes at room temperature followed by washing in FACS buffer. Cells were then incubated with Fc block (1:500 dilution, Cat. # 553142, clone 2.4G2, BD Biosciences) diluted in FACS buffer for 5 minutes to block nonspecific binding of antibodies to cellular Fc receptors. Cells were then incubated with fluorochrome-conjugated antibodies in various combinations to allow for multi-staining for 30 minutes at 4°C. Cells were washed three times with FACS buffer and fixed in 200 µl of 2% ultrapure formaldehyde (Polysciences, Warrington, PA) diluted in FACS buffer (fixation buffer). Fluorescence minus one (FMO) controls or cells incubated with either FACS buffer alone, or single fluorochrome-conjugated antibodies were used to determine positive staining and spillover/compensation calculations, and background fluorescence was determined with FlowJo v.10.8 Software (FlowJo, LLC, Ashland, OR). Raw data were collected with a Cell Analyzer LSRII (BD Biosciences) using BD FACSDiva v8.0 software at the University of North Texas Health Sciences Center (UNTHSC) Flow Core, and compensation and data analyses were performed using FlowJo v.10.8 Software. Cells were first gated for lymphocytes (SSC-A vs. FSC-A) and singlets (FSC-H vs. FSC-A). The singlets gate was further analyzed for the uptake of live/dead yellow stain to determine live vs. dead cells. From live cells, cells were gated on CD45+ cell expression. For data analyses, 100,000 events (cells) were evaluated from a predominantly leukocyte population identified by back gating from CD45+ stained cells. Statistical significance between strains at each timepoint was determined using Student’s *t* test (GraphPad Software, San Diego, CA).

### Macrophage activation analyses

Intracellular staining of markers of macrophage activation was performed as described previously (29). Leukocytes isolated from infected mice as described above were incubated with cell stimulation cocktail (eBioscience Cat. # 00-4970-03) according to the manufacturer’s recommendation and incubated at 37°C in 5% CO_2_ in cRPMI for two hours in a six-well plate. Golgi plug (1:100 dilution, Brefeldin A, Cat. # 51-2301KZ, BD Biosciences) was added according to the manufacturer’s recommendations and incubated for an additional four hours (6 hours total). Cells were washed with PBS and stained with yellow Zombie viability dye in PBS at room temperature in the dark for 15 minutes. Cells were then washed with FACS buffer and incubated with Fc block (BD Biosciences) diluted in FACS buffer for 5 minutes. For nitric oxide (iNOS) and Arginase 1 (Arg1) production in macrophages, cells were stained for surface markers CD45, CD11b, CD64, F4/80, and CD24, and incubated at 4°C for 30 minutes. Cells were then washed and fixed with 2% ultra-pure formaldehyde (Polysciences, Warrington, PA) for 20 minutes. Subsequently, cells were washed with 0.1% saponin buffer and stained with antibodies for iNOS and Arg1 for 30 minutes at 4°C. Finally, cells were washed with saponin buffer and fixed with 2% ultra-pure formaldehyde. Samples were processed using a Cell Analyzer LSRII (BD Biosciences) using BD FACSDiva v8.0 software at the UNTHSC Flow Core, and 100,000 events were collected for analysis using FlowJo v.10.8 Software. Statistical significance between strains at each timepoint was determined using Student’s *t* test (GraphPad Software, San Diego, CA).

### Titan cell assay and quantification

A previously described *in vitro* titanization assay was used here (30). In brief, the WT (H99), the *mar1*Δ mutant (MAK1), and the *mar1*Δ + *MAR1* (MAK11) strains were incubated for 18 hours at 30°C, 150 rpm in 5 mL yeast nitrogen base (YNB) without amino acids prepared according to the manufacturer’s instructions plus 2% glucose. Cultures were washed six times with PBS. An optical density at 600 nm (OD_600_) of 0.001 for each strain was transferred to 5 mL 10% heat-inactivated fetal bovine serum (HI-FBS) in PBS and incubated at 37°C, 5% CO_2_ for 96 hours. Cells were imaged by differential interference contrast (DIC) microscopy using a Zeiss Axio Imager A1 microscope equipped with an Axio-Cam MRm digital camera. Cell diameter was measured using the ImageJ software (FIJI), and cells with a diameter > 10 μm were considered Titan cells. A minimum of 400 cells were analyzed across three biological replicates for each fungal strain. Statistical significance was determined using one-way analysis of variance (ANOVA) and the Tukey-Kramer test (GraphPad Software, San Diego, CA).

### SEM polysaccharide capsule visualization

The WT (H99), the *mar1*Δ mutant (MAK1), the *mar1*Δ + *MAR1* (MAK11), and the *cap59*Δ mutant (cap59) strains were incubated in YPD medium at 30°C and CO_2_-independent medium (Gibco) at 37°C until saturation. Samples were fixed with 2.5% glutaraldehyde for 1 hour at room temperature and were subsequently washed 3 times with PBS. Each sample was mounted onto 12 mm poly-L-lysine-coated coverslips (Neuvitro Corporation) and subsequently dehydrated by immersing the coverslips in ethanol (30% for 5 minutes, 50% for 5 minutes, 70% for 5 minutes, 95% for 10 minutes, 100% for 10 minutes, and 100% for 10 minutes). Samples were then critical point dried with a Tousimis 931 critical point dryer (Rockville, Maryland) and coated with gold-palladium using a Cressington 108 sputter-coater (Watford, United Kingdom). Coverslips containing the prepared samples were mounted and imaged on a Hitachi S-4700 scanning electron microscope (Tokyo, Japan).

### Cellular morphology defect quantification

The WT (H99), the *mar1*Δ mutant (MAK1), and the *mar1*Δ + *MAR1* (MAK11) strains were incubated for 18 hours in YPD medium at 30°C with shaking at 150 rpm. An OD_600_ of approximately 0.2 for each strain was transferred to fresh YPD medium and subsequently incubated at either 30°C or 37°C for 18 hours with shaking at 150 rpm. Cells were then pelleted, washed with PBS, and imaged by differential interference contrast (DIC) microscopy. DIC images were captured using a Zeiss Axio Imager A1 microscope equipped with an Axio-Cam MRm digital camera. A minimum of 500 cells were analyzed across three biological replicates for each strain using the ImageJ software (FIJI). Statistical significance was determined using two-way analysis of variance (ANOVA) and the Tukey-Kramer test (GraphPad Software, San Diego, CA).

### Growth curve analysis

The WT (H99), the *mar1*Δ mutant (MAK1), and the *mar1*Δ + *MAR1* (MAK11) strains were incubated for 18 hours in YPD medium at 30°C with 150 rpm shaking. Cultures were normalized to an OD_600_ of 0.01 in fresh YPD medium and added to wells of a 96-well plate. Growth was then measured at an absorbance of 595 nm every 10 minutes for 40 hours with shaking between readings and incubation at 37°C. Control wells containing YPD medium alone were also included to eliminate any background absorbance.

### Hypoxia resistance analyses

The WT (H99), the *mar1*Δ mutant (MAK1), the *mar1*Δ + *MAR1* (MAK11), and the *sre1*Δ mutant (HEB6) strains were incubated in YPD medium at 30°C until mid-logarithmic growth phase. Strains were washed once in PBS, normalized to an OD_600_ of 0.6 in PBS, and serially diluted onto YES (0.5% [w/v] yeast extract, 2% glucose, and 225 µg/mL uracil, adenine, leucine, histidine, and lysine) medium agar plates with or without cobalt chloride (0.7 mM) (31). Microaerophilic conditions were generated using System sachets (31). Plates were placed in the chamber (microaerophilic) or outside the chamber (ambient air), incubated at 30°C, and imaged daily for 96 hours.

### Mouse isolate recovery and phenotypic characterization

C57BL/6 female mice acquired from Charles River Laboratories were infected as described above. At 61 DPI and 100 DPI, mice were sacrificed by CO_2_ inhalation followed by an approved secondary method of euthanasia and fungi were subsequently isolated from the lungs as described above. Single fungal colonies were plated onto YPD agar medium and subsequently frozen in separate wells of 96-well plates at -80°C. Isolated fungi were stamped onto YPD agar medium incubated at 30°C, YPD agar medium incubated at 37°C, YPD agar medium supplemented with nourseothricin (NAT) (100 µg/mL) incubated at 30°C, and YPD agar medium buffered (150 mM HEPES) to pH 8.15 incubated at 30°C. All plates were imaged daily. Mouse isolates were determined to be *mar1*Δ mutant strain isolates based on growth on YPD + NAT medium and dry colony morphology on YPD pH 8.15 medium (23). The original WT (H99) and *mar1*Δ mutant (MAK1) strains were included on each plate as controls.

### Ethical use of animals

All animal experiments in this manuscript were approved by the University of Texas at San Antonio Institutional Animal Care and Use Committee (IACUC) (protocol #MU021), the Texas Christian University and the University of North Texas Health Sciences Center (UNTHSC) IACUC (protocol #1920-9), and the Duke University IACUC (protocol #A102-20-05). Mice were handled according to IACUC guidelines.

## Data availability

All fungal strains and reagents are available upon request.

## RESULTS

### Pulmonary granulomas are formed and maintained in mice infected with the *mar1*Δ mutant strain

Based on our recent observations that the *mar1*Δ mutant strain displays a highly immunogenic cell surface, we hypothesized that the *mar1*Δ mutant strain would have unique interactions with the host *in vivo*. We previously observed that the *mar1*Δ mutant strain is hypovirulent compared to the wild-type (WT) strain in a murine inhalation model of cryptococcosis (23). Highly immunogenic fungal strains often induce a hyperinflammatory response that is detrimental to the host, resulting in hypervirulence (32–34). We therefore explored in greater detail the mechanisms by which the highly immunogenic *mar1*Δ mutant strain simultaneously activates and is controlled by the host immune response.

As an initial investigation into the interactions between the *mar1*Δ mutant strain and the host, we assessed the gross appearance of infected lungs from our previously reported *mar1*Δ mutant strain murine inhalation infection experiment. At the time of sacrifice, generally between 24-40 days post-inoculation (DPI), we observed that the lungs of *mar1*Δ-infected C57BL/6 mice displayed large, well-circumscribed inflammatory foci surrounded by healthy-appearing lung tissue (Figure 1A). This contrasts starkly with WT-infected lungs, which typically exhibit uncontrolled fungal proliferation accompanied by a diffuse inflammatory response.

**Figure 1.**
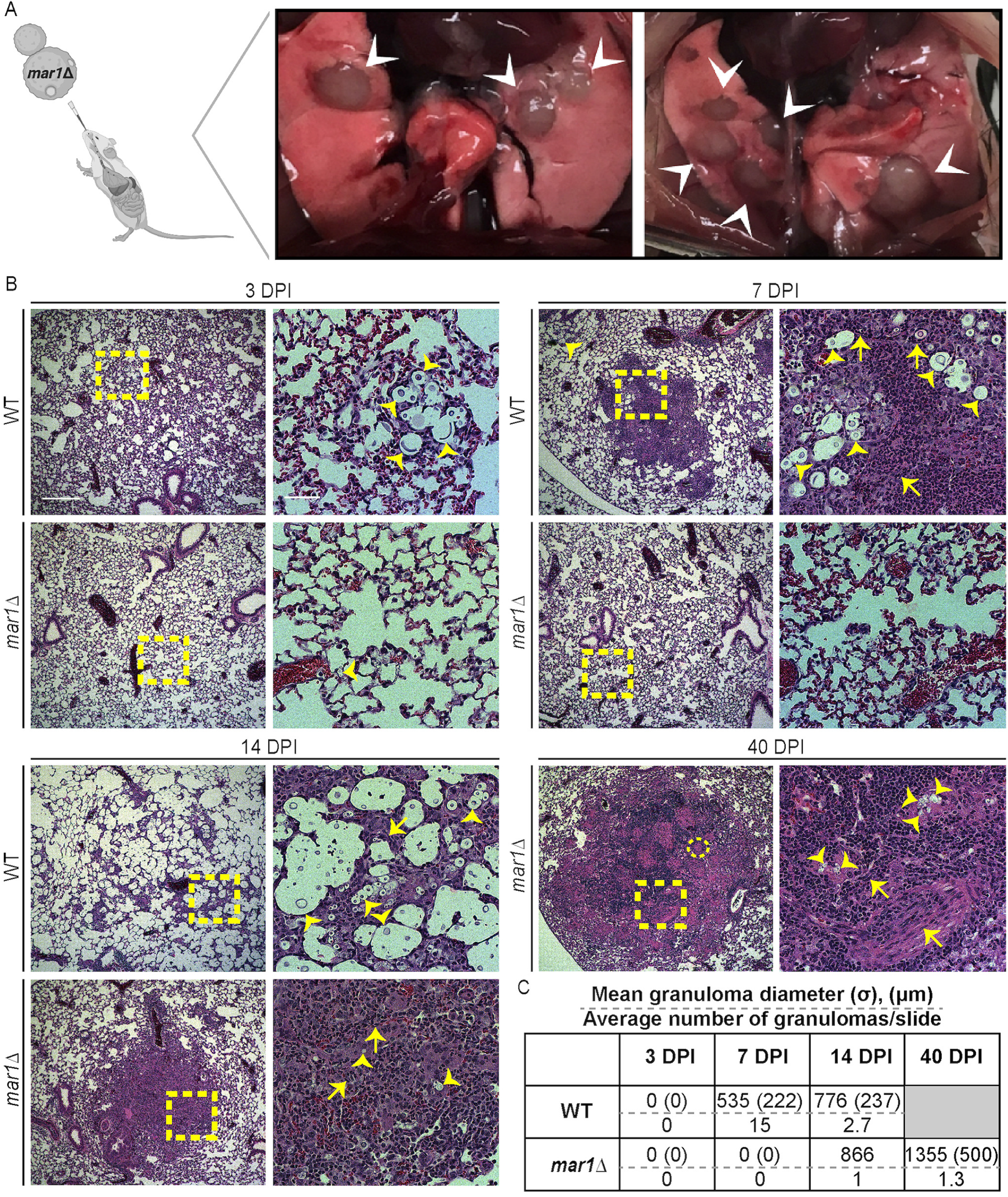
Pulmonary granuloma formation in murine cryptococcal infections. A. Lung dissections of female C57BL/6 mice infected with the *mar1*Δ mutant strain were performed to display macroscopic lung pathology, specifically granulomas (white arrowheads). Cartoon adapted from BioRender.com (2021). B. The lungs of female C57BL/6 mice inoculated with 1 × 10^4^ cells of the WT strain or the *mar1*Δ mutant strain sacrificed at predetermined endpoints (3, 7, 14, and 40 DPI) were harvested for histopathological analyses. Hematoxylin and eosin staining were utilized to visualize microscopic lung pathology (fungal cells [yellow arrowheads], multinucleated giant cells [yellow circle], epithelioid macrophages (yellow arrows), inset [yellow boxes]). 5x scale bar (left), 250 μm. 10x scale bar (right), 50 μm. C. Granuloma diameter (μm) was measured using FIJI. σ, standard deviation (μm). Gray box, no experimental subjects can be assessed at this timepoint.

We examined histopathological features of infected murine lungs at specific timepoints throughout the course of infection to further characterize the unique pathology observed in *mar1*Δ-infected lungs. To do so, we replicated the experimental approach used in Figure 1A; we inoculated C57BL/6 mice by inhalation with the WT strain or the *mar1*Δ mutant strain and subsequently harvested lungs for analysis throughout infection. At 3 DPI, an early timepoint in infection at which all inoculated mice still appear healthy, the WT-infected and *mar1*Δ-infected lungs appear similar, with the only notable exception being the increased number of fungal cells observed in the WT-infected lungs (Figures 1B & S1). By 7 DPI, a timepoint in infection in which WT-infected mice begin to show signs and symptoms of fungal disease but the *mar1*Δ-infected mice still appear healthy, WT-infected lungs display numerous small foci of inflammation (mean diameter = 535 μm) that contain some, but not all, fungal cells (Figures 1B, 1C, & S1). As described previously, many of the WT fungal cells exhibit signs of titanization (35). These foci of inflammation also occasionally display hallmarks of early granuloma formation, such as the presence of epithelioid macrophages (1, 2, 4) (Figure 1B). This type of immature granulomatous inflammatory response has been reported previously in the C57BL/6 background infected with the *C. neoformans* serotype D strain, 52D (17). In contrast, the *mar1*Δ-infected lungs have few visible fungal cells and display a more uniform pattern of inflammation throughout the lungs at 7 DPI (Figures 1B). These observations demonstrate that distinct characteristics of the *mar1*Δ mutant strain pathology emerge early in infection.

At 14 DPI, a timepoint in infection in which the WT-infected mice begin to succumb to fungal infection and the *mar1*Δ-infected mice still appear healthy, WT cells, many of which are titanized, proliferate throughout the lungs resulting in a scattered, unorganized inflammatory response with mixed cell infiltrates (Figure 1B). Additionally, most nascent granulomas have broken down, which may explain why this timepoint also corresponds to a period of accelerating clinical symptoms and imminent mortality in WT-infected mice (Figures 1B & 1C). In contrast, *mar1*Δ-infected lungs begin to form granulomas by 14 DPI. Specifically, foci of inflammation appear (mean diameter = 866 μm), containing few fungal cells which are surrounded by regions of normal-appearing lung tissue without fungal or inflammatory cells (Figures 1B & 1C). Additionally, these inflammatory foci contain hallmarks of granulomas, such as epithelioid macrophages surrounded by lymphocytes (1, 2, 4) (Figure 1B). In contrast to those infected with the WT strain, the *mar1*Δ-infected mice display few infection-related symptoms at this timepoint.

Many *mar1*Δ-infected mice survive to 40 DPI (23). At this late timepoint in infection, mature granulomas are frequently observed (mean diameter = 1355 μm), containing fungal cells, multinucleated giant cells, and palisading epithelioid macrophages (Figures 1B, 1C, & S1). Additionally, no fungal cells are observed in lung tissue outside of these granulomas. Collectively, these observations suggest differences in the immune response in the context of WT and *mar1*Δ mutant strain infection. WT-infected mice show a consistently robust mixed inflammatory response and variable Titan cell response with vague granuloma formation during early stages of infection (7 DPI) (Figures 1B, 1C, & S1). This response is ineffective and is quickly overcome by fungal growth, resulting in fungal proliferation throughout the lungs (14 DPI) (Figures 1B & 1C). In contrast, the *mar1*Δ-infected mice show an absent to minimal inflammatory response, absent Titan cell formation, and minimal granulomatous inflammatory response during early stages of infection (7 DPI), with a more well-formed granulomatous response in mice that survive to later timepoints in infection (40 DPI) (Figures 1B, 1C, & S1). These *mar1*Δ-induced granulomas appear sufficient to contain fungal proliferation.

### The *mar1*Δ mutant strain has a reduced fungal burden and hyper-immunogenicity *in vivo*

To explore possible mechanisms by which *mar1*Δ mutant strain infections induce pulmonary granuloma formation, we assessed fungal burden and the pulmonary immune response at timepoints relevant to granuloma formation. To do so, we replicated the experimental approaches used in Figure 1; we inoculated C57BL/6 mice by inhalation and harvested lungs for analysis throughout infection. We previously reported a decrease in fungal burden in *mar1*Δ-infected lungs compared to WT-infected lungs as early as 1 and 4 DPI, despite identical doses being used for both strains (23). In this work, at all tested timepoints (3, 7, 14, & 21 DPI), we find that *mar1*Δ-infected lungs have a significantly reduced fungal burden compared to WT-infected lungs. Specifically, the *mar1*Δ-infected lungs have a 10-fold reduction in fungal burden at 3 DPI, a 100-fold reduction in fungal burden at 7 DPI, and a >500-fold reduction in fungal burden at 14 and 21 DPI compared to WT-infected lungs (Figure 2). These observations support the reduced number of *mar1*Δ mutant cells observed at these same timepoints in our histopathology analyses (Figure 1B). As a result of the drastic reduction in pulmonary fungal burden throughout infection, we observed that the *mar1*Δ mutant strain rarely disseminates to the brain (Figure 2). When the *mar1*Δ mutant strain does disseminate to the brain, the fungal burden is markedly lower than that of the WT strain (Figure 2). Together, these observations indicate that the *mar1*Δ mutant strain has reduced fungal burden in the murine lung and brain, reinforcing our previous reports that the *mar1*Δ mutant strain has reduced fitness in host-relevant conditions.

**Figure 2.**
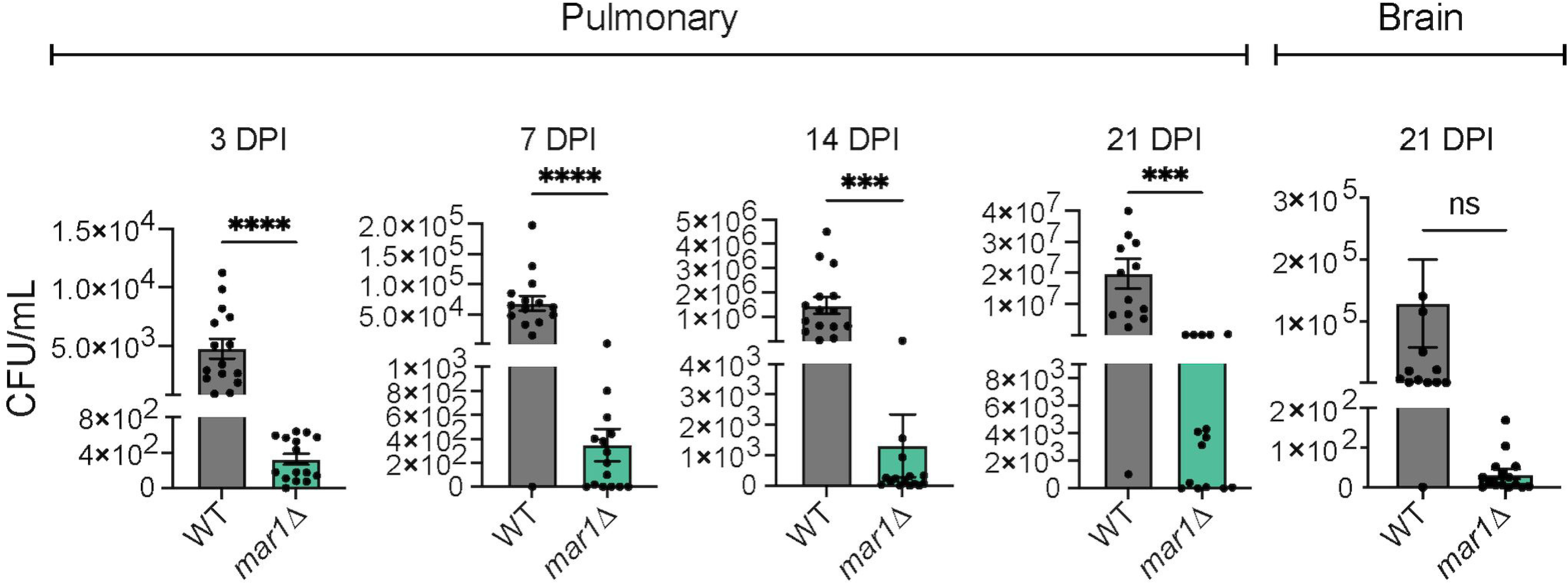
Fungal burden throughout infection. Pulmonary fungal burden of female C57BL/6 mice (*n* = 15) inoculated with 1 × 10^4^ cells of the WT strain or the *mar1*Δ mutant strain was measured by quantitative cultures throughout infection: 3, 7, 14, and 21 DPI. Brain fungal burden of female C57BL/6 mice (*n* = 15) inoculated with 1 × 10^4^ cells of the WT strain or the *mar1*Δ mutant strain was measured by quantitative cultures at 21 DPI. Error bars represent standard error of the mean (SEM). Statistical significance was determined using Student’s *t* test (***, *P* < 0.001; ****, *P* < 0.0001; ns, not significant).

Based on the drastic differences in fungal burden observed between WT-infected and *mar1*Δ-infected lungs, we hypothesized that the immune microenvironment within the lungs would also differ significantly. We replicated the experimental approaches used in Figure 1; we inoculated C57BL/6 mice by inhalation and harvested lungs for analysis throughout infection. At early timepoints in infection, 1 and 3 DPI, we observed similar pulmonary cytokine profiles and leukocyte infiltrates within WT-infected lungs and *mar1*Δ-infected lungs (Figures 3A, 3B, S2, & S3). The only significant difference observed between the two infections was in the production of granulocyte macrophage-colony stimulating factor (GM-CSF), a cytokine required for maturation of myeloid cells. Specifically, *mar1*Δ-infected lungs display a significant increase in GM-CSF production compared to WT-infected lungs at 3 DPI, an early timepoint in infection at which the pulmonary immune response is being actively developed (Figures 3A & S2). Despite the drastic reduction in *mar1*Δ mutant fungal burden at these early timepoints, the *mar1*Δ mutant strain induces a cytokine and cellular response comparable to that of the WT strain, likely due to the increased immunogenicity of the *mar1*Δ mutant cells.

**Figure 3.**
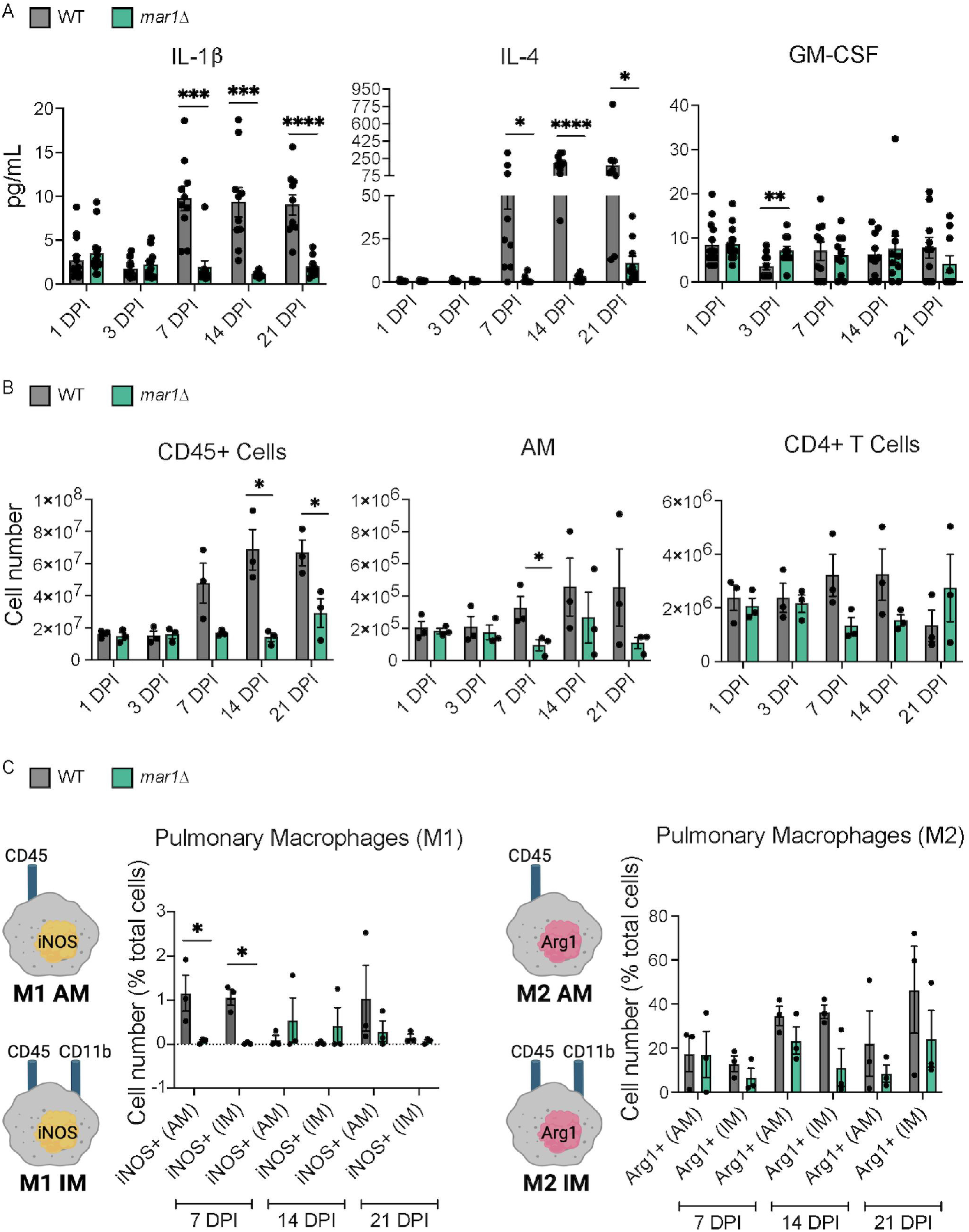
Pulmonary cytokine profile and leukocyte infiltrate associated with granuloma formation. A. Pulmonary cytokine responses of female C57BL/6 mice inoculated with 1 × 10^4^ cells of the WT strain or the *mar1*Δ mutant strain were measured using the Bio-Plex protein array system throughout infection: 1 (*n* = 15), 3 (*n* = 15), 7 (*n* = 10), 14 (*n* = 10), and 21 (*n* = 10) DPI. Error bars represent SEM. Statistical significance between strains at each timepoint was determined using Student’s *t* test (*, *P* < 0.05; **, *P* < 0.01; ***, *P* < 0.001; ****, *P* < 0.0001; no designation between strains, not significant). Only a subset of data is shown; refer to Figure S2 for full analysis. B. Pulmonary leukocyte infiltrates of female C57BL/6 mice inoculated with 1 × 10^4^ cells of the WT strain or the *mar1*Δ mutant strain were measured by flow cytometry throughout infection: 1, 3, 7, and 21 DPI. Data shown are the mean ± of absolute cell numbers from three independent experiments (*n* = 3) performed using five mice per group per timepoint per experiment. Error bars represent SEM. Statistical significance between strains at each timepoint was determined using Student’s *t* test (*, *P* < 0.05; no designation between strains, not significant). Only a subset of data is shown; refer to Figure S3 for full analysis. C. Pulmonary macrophage activation of female C57BL/6 mice (*n* = 3) inoculated with 1 × 10^4^ cells of the WT strain or the *mar1*Δ mutant strain were measured by flow cytometry throughout infection: 7, 14, and 21 DPI. Inducible nitrogen oxide synthase (iNOS) was used as a marker for M1 macrophages and Arginase 1 (Arg1) was used as a marker for M2 macrophages. The percentage of total iNOS+ cells and Arg1+ cells is shown. Error bars represent the SEM. Log transformation was used to normally distribute the data for statistical analysis. Statistical significance between strains at each timepoint was determined using Student’s *t* test (*, *P* < 0.05; no designation between strains, not significant). AM = alveolar macrophage (CD45+, CD11b-). IM = interstitial macrophage (CD45+, CD11b+). Cartoons adapted from BioRender.com (2021).

As infection progressed to 7, 14, and 21 DPI, we observed marked reductions in multiple cytokines (including IL-1β, IL-4, and GM-CSF) and leukocytes (including CD45+ cells, alveolar macrophages [AM], and CD4+ T cells) in *mar1*Δ-infected lungs compared to WT-infected lungs (Figures 3A, 3B, S2, & S3). These observations demonstrate that by these timepoints in infection, the overall cytokine and cellular response is reduced in *mar1*Δ-infected lungs compared to WT-infected lungs, likely due to the sustained reduction in fungal burden present in the *mar1*Δ-infected lungs. This is further supported by our histopathological observations made at the same timepoints demonstrating more localized regions of inflammation in *mar1*Δ-infected lungs than in WT-infected lungs (Figure 1B). We further explored macrophage polarization at these same timepoints to determine whether the reduction in *mar1*Δ mutant strain fungal burden, and the subsequent reduction in the pulmonary immune response, are due to differences in macrophage activation (36). At each tested timepoint (7, 14, and 21 DPI), we observed that the *mar1*Δ-infected lungs have a comparable number of or fewer classically-activated (M1) and alternatively-activated (M2) alveolar and interstitial macrophages compared to WT-infected lungs (Figure 3C). These observations demonstrate that the *mar1*Δ mutant strain does not induce differential macrophage polarization that results in a reduction in fungal burden and a more protective immune response. Collectively, these data suggest that *mar1*Δ-induced pulmonary granuloma formation appears to be a largely fungal-driven phenomenon. Despite reductions in fungal burden, the *mar1*Δ mutant strain induces a WT strain-like immune response early in infection that results in fungal containment within granulomas during mid-late stages of infection. As infection matures and progresses, there is a marked decrease in many cytokines and leukocytes infiltrating the *mar1*Δ-infected lung that corresponds with the sustained reduction in fungal burden.

### Host GM-CSF signaling is required for pulmonary granuloma formation

Granuloma formation is dependent on GM-CSF signaling in the context of both mycobacterial (24–26) and cryptococcal infections (17). GM-CSF is the only cytokine that showed significant differential production in our cytokine analyses. Specifically, we observed that the *mar1*Δ mutant strain induces more pulmonary GM-CSF production than the WT strain at 3 DPI (Figures 3A & S2). We therefore hypothesized that GM-CSF signaling would also be required for the formation of pulmonary granulomas in our model. To test this hypothesis, we assessed the progression of infections with the WT strain and the *mar1*Δ mutant strain in the Csf2rb^-/-^ mouse background, which is defective in GM-CSF signaling due to loss of the functional GM-CSF receptor. We inoculated Csf2rb^-/-^ mice using the inhalation route and harvested lungs for analysis throughout infection. Overall, a similar pattern of inflammation was observed between mice infected with the WT strain and mice infected with the *mar1*Δ mutant strain. We observed that putative pulmonary granulomas are absent in Csf2rb^-/-^ mice infected with either strain at every tested timepoint (3, 7, and 14 DPI) (Figures 4A & S1). Instead, inflammation appears unorganized and diffuse throughout the entirety of the lungs infected with either fungal strain. Contrasting with the C57BL/6 infections, the Csf2rb^-/-^ infections appear to be characterized by fewer macrophages, which is expected based on previous work that demonstrated that GM-CSF is required for macrophage recruitment to the lung during early cryptococcal infection (17) (Figures 4A & S1). Like the C57BL/6 infections, however, WT fungal cells are abundant throughout the lung, many with signs of titanization, while *mar1*Δ mutant fungal cells are infrequently observed (Figures 4A & S1). Pulmonary fungal burden assessed at 3 DPI confirms that *mar1*Δ-infected lungs have a significantly lower fungal burden, with a 10-fold reduction compared to WT-infected lungs, similar to what was observed in the C57BL/6 infections (Figure 4B). These data demonstrate that GM-CSF signaling is required for granuloma formation in both WT strain and *mar1*Δ mutant strain infections. However, because loss of GM-CSF signaling does not rescue the reduction of *mar1*Δ mutant strain fungal burden during early stages of infection, these data also suggest that GM-CSF signaling does not exclusively drive the impaired fitness of *mar1*Δ mutant cells in the murine lung.

**Figure 4.**
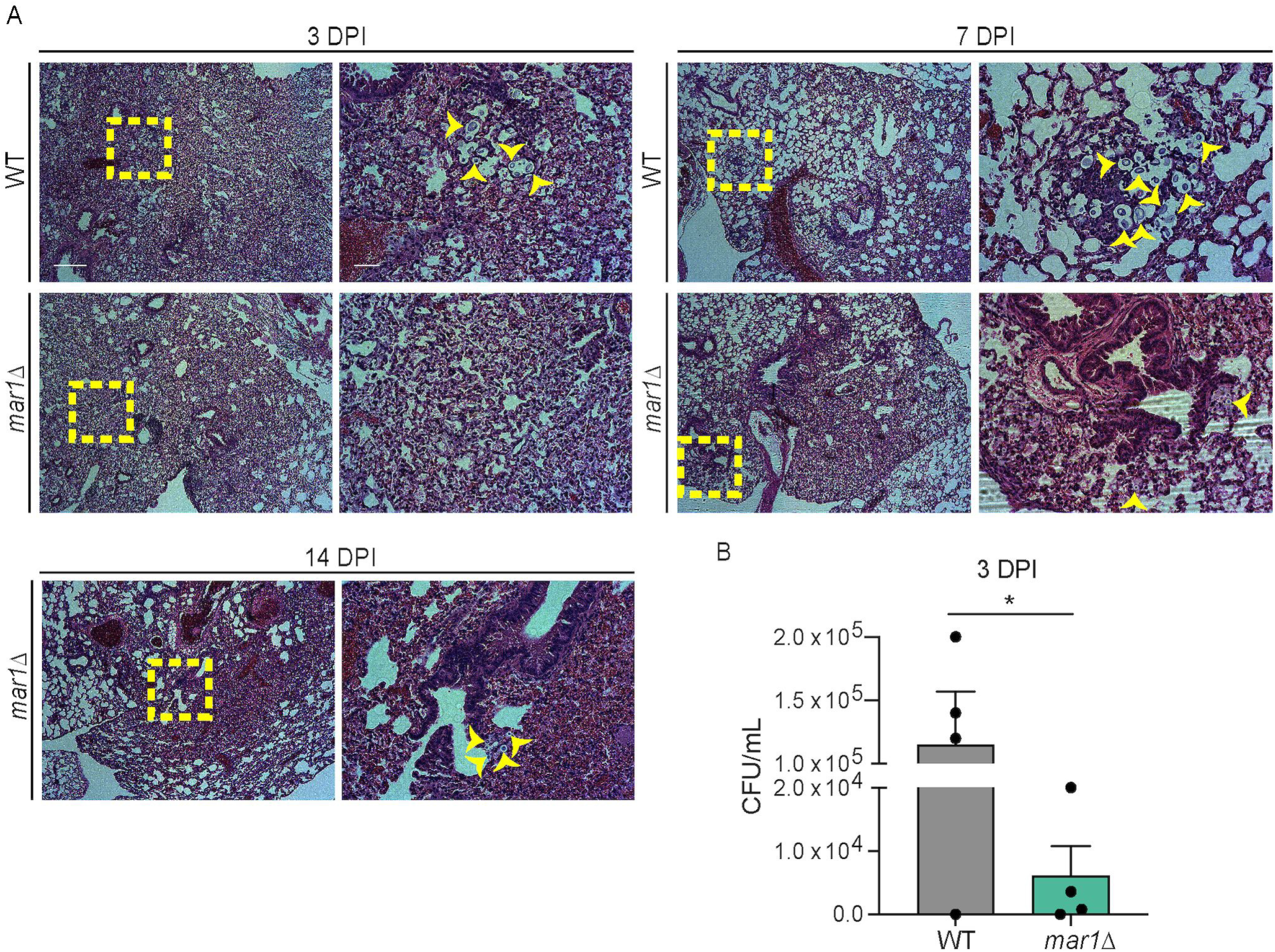
Contributions of GM-CSF signaling to pulmonary granuloma formation. A. The lungs of female (*n* = 2) (shown) and male (*n* = 2) (not shown) Csf2rb^-/-^ mice inoculated with 1 × 10^4^ cells of the WT strain or the *mar1*Δ mutant strain sacrificed at predetermined endpoints (3, 7, and 14 DPI) were harvested for histopathological analyses. Hematoxylin and eosin staining were utilized to visualize microscopic lung pathology (fungal cells [yellow arrowheads], inset [yellow boxes]). 5x scale bar (left), 250 μm. 10x scale bar (right), 50 μm. B. Pulmonary fungal burden of female (*n* = 2) and male (*n* = 2) Csf2rb^-/-^ mice inoculated with 1 × 10^4^ cells of the WT strain or the *mar1*Δ mutant strain sacrificed at 3 DPI was measured by quantitative cultures. Error bars represent the SEM. Statistical significance was determined using Student’s *t* test (*, *P* < 0.05).

### The *mar1*Δ mutant strain is attenuated in the employment of various virulence factors

In our fungal burden assays, we observed a modest increase in *mar1*Δ mutant strain fungal burden as infection progressed from 3 to 21 DPI (Figure 2). Despite this, we find that *mar1*Δ-infected mice can remain healthy-appearing and survive to at least 100 DPI. Furthermore, viable *mar1*Δ mutant cells that retain previously reported *mar1*Δ mutant phenotypes, including dry colony morphology on alkaline pH and nourseothricin (NAT) resistance (23), can be recovered from the lung at extended timepoints in infection (61 and 100 DPI) (Figure S4). These observations indicate that the *mar1*Δ mutant strain can persist within murine lung granulomas for extended periods of time without causing any symptoms or signs of disease. Based on this observation, we sought to understand the mechanism by which the *mar1*Δ mutant strain can survive and persist in the mouse lung.

In both human and murine infections, a subset of cryptococcal cells form enlarged Titan cells, an important virulence factor that enables cryptococcal persistence in the lungs (35, 37). Using an established *in vitro* titanization assay (30), we observed that the *mar1*Δ mutant strain is unable to form Titan cells (Figure 5A). This observation supports our histopathology experiments, in which Titan cells were absent in *mar1*Δ-infected lungs, collectively demonstrating that Titan cell formation does not explain the persistence of *mar1*Δ mutant cells within granulomas.

**Figure 5.**
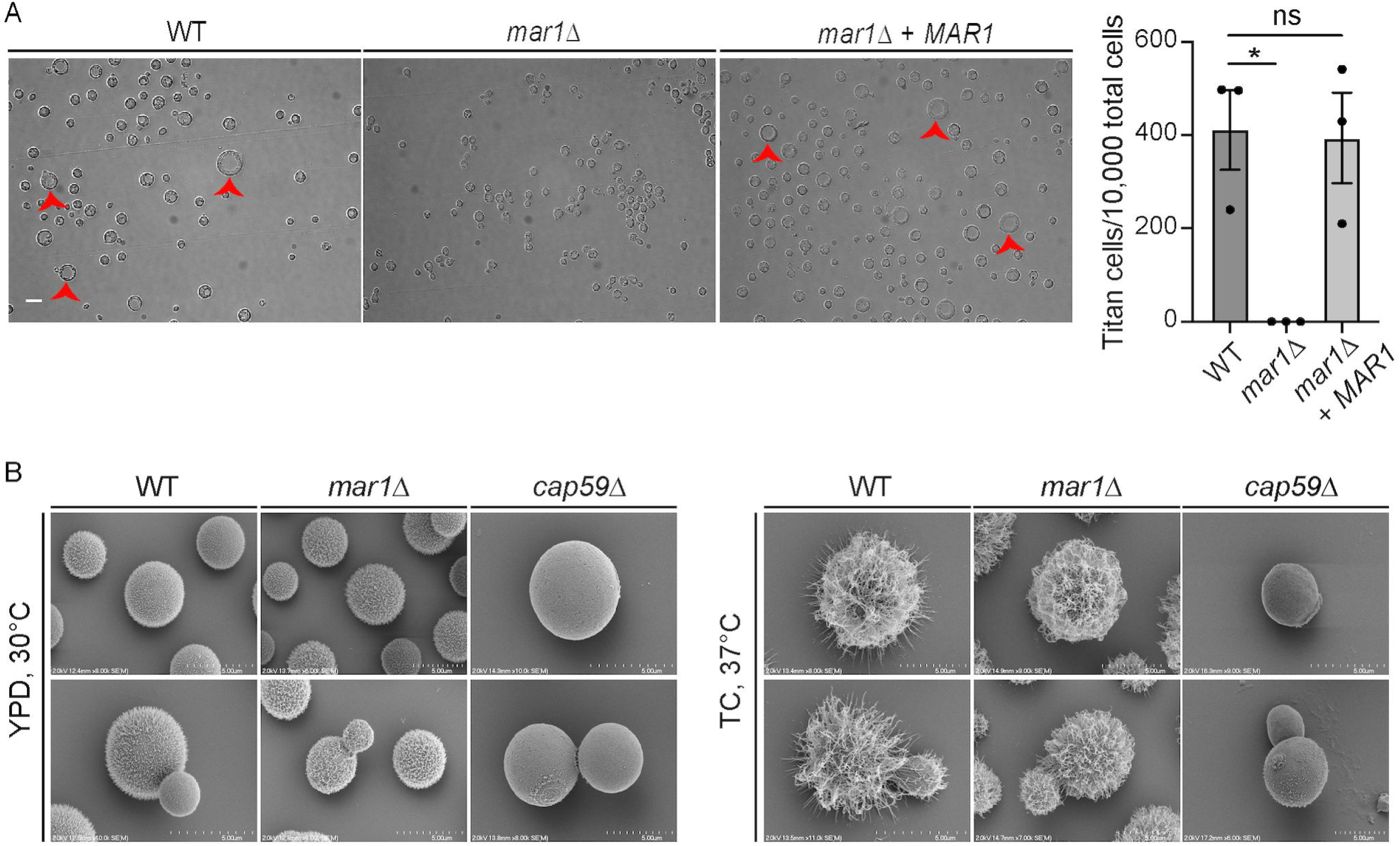
Cell cycle-mediated virulence factor phenotypes of the *mar1*Δ mutant strain. A. Titan cell formation was induced in the WT strain, the *mar1*Δ mutant strain, and the *mar1*Δ + *MAR1* complemented strain. Cells were pre-grown in YNB medium at 30°C and an OD_600_ of 0.001 was transferred to 10% HI-FBS in PBS incubated at 5% CO_2_, 37°C for 96 hours. Cells were imaged by DIC microscopy (Zeiss Axio Imager A1). Cell diameter was measured using FIJI, and cells with a diameter > 10 μm were considered Titan cells (red arrowheads). The number of Titan cells per 10,000 cells was calculated for each strain. A minimum of 400 cells were analyzed across three biological replicates (*n* = 3). Error bars represent the SEM. Statistical significance was determined using a one-way ANOVA (*, *P* < 0.05; ns, not significant). 63x scale bar, 10 μm. B. The WT strain, the *mar1*Δ mutant strain, the *mar1*Δ + *MAR1* complemented strain, and the *cap59*Δ mutant strain were incubated in YPD medium at 30°C and CO_2_-independent medium (TC) at 37°C until saturation. Samples were subsequently fixed, mounted, dehydrated, and sputter-coated. Samples were imaged with a Hitachi S-4700 scanning electron microscope to visualize capsule organization and elaboration.

We previously reported that the *mar1*Δ mutant strain is impaired in the implementation of the polysaccharide capsule, assessed by India ink staining (23). We utilized high-resolution scanning electron microscopy (SEM) to more rigorously study the *mar1*Δ mutant strain capsule architecture. In permissive growth conditions (YPD medium, 30°C), the capsule of the *mar1*Δ mutant strain is nearly indistinguishable from that of the WT strain, which contrasts starkly with the acapsular *cap59*Δ mutant strain (Figure 5B). However, in capsule-inducing conditions (TC medium, 37°C), the *mar1*Δ mutant strain lacks the degree of capsule fiber elongation observed in the WT strain, explaining the reduction in India ink exclusion previously reported for the *mar1*Δ mutant strain (23) (Figure 5B). The inability of the *mar1*Δ mutant strain to employ these two important virulence factors, Titan cells and polysaccharide capsule, likely drive the hyper-immunogenicity observed in our murine infection studies.

### The *mar1*Δ mutant strain displays cell cycle defects that result in a slow growth phenotype and hypoxia resistance

Both Titan cell formation (30, 38) and polysaccharide capsule elaboration (39–41) are known to be mediated by the cell cycle, suggesting that the *mar1*Δ mutant strain may be unable to properly employ these virulence factors due to defects in cell cycle progression. To explore cell cycle progression in the *mar1*Δ mutant strain background, we observed *mar1*Δ mutant cell morphology during logarithmic growth phase. When incubated at the permissive temperature of 30°C, the *mar1*Δ mutant strain displays an increased incidence of cytokinesis defects (such as elongated cells, cells with wide bud necks, and cells that fail to complete cytokinesis), compared to both the WT strain and the *mar1*Δ + *MAR1* complemented strain (Figure 6A). The frequency of these cytokinesis defects is significantly enhanced at the physiological temperature of 37°C (Figure 6A). We next determined the impact of these defects on the growth kinetics of the *mar1*Δ mutant strain. We observed that the *mar1*Δ mutant strain displays a reduction in growth during logarithmic phase at 37°C, compared to both the WT strain and the *mar1*Δ + *MAR1* complemented strain (Figure 6B). These data demonstrate that the *mar1*Δ mutant strain has a slow growth phenotype at the physiological temperature of 37°C that is likely driven in part by cytokinesis defects.

**Figure 6.**
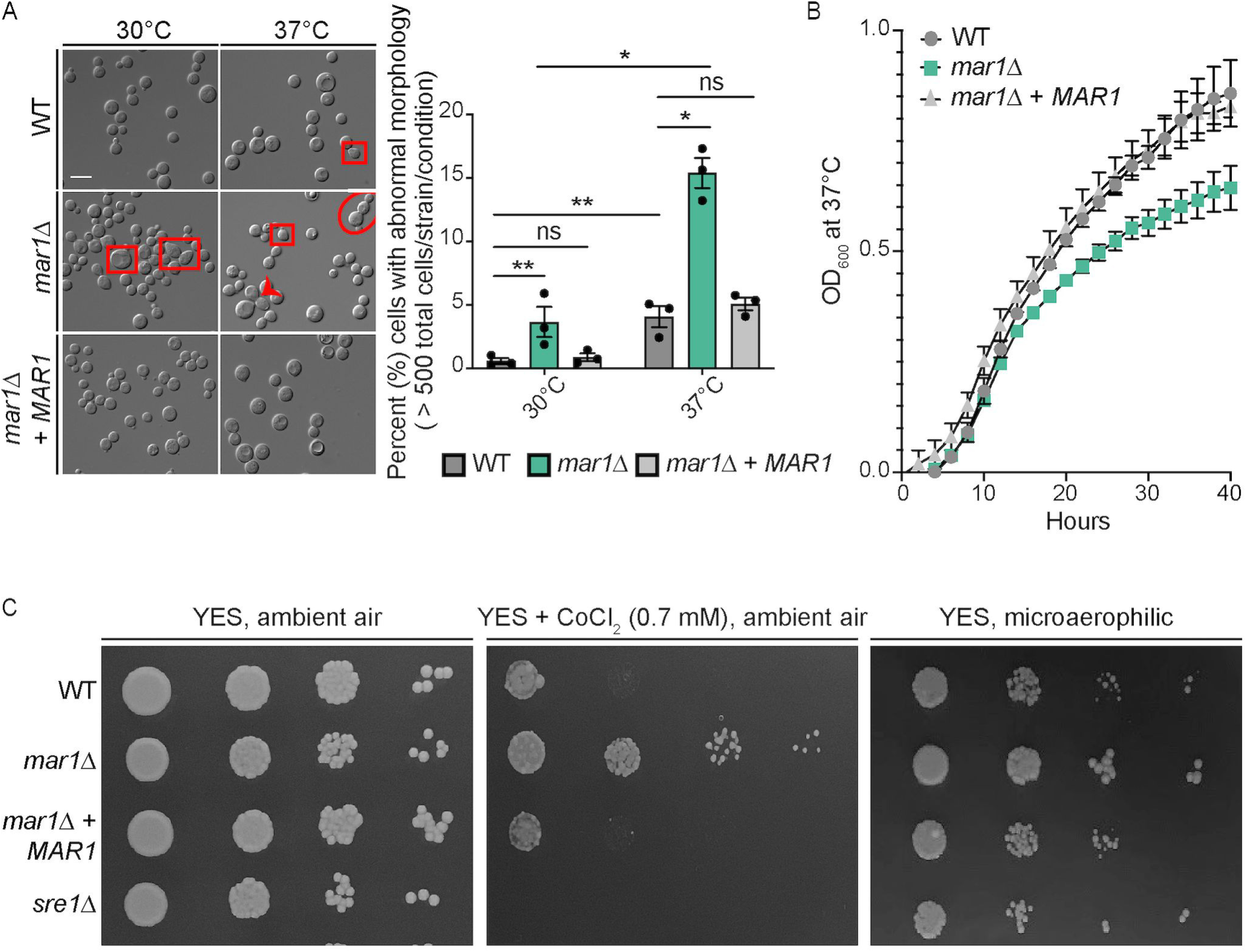
Slow growth phenotypes of the *mar1*Δ mutant strain. A. Morphological defects were analyzed in the WT strain, the *mar1*Δ mutant strain, and the *mar1*Δ + *MAR1* complemented strain through incubation in YPD medium at either 30°C or 37°C. Cells were imaged by DIC microscopy (Zeiss Axio Imager A1) and were subsequently visually inspected for morphological defects, such as elongated cells (red squares), wide bud necks (red arrowhead), and cytokinesis failure (red circle). The percentage of total cells displaying morphological defects was quantified for each strain at each temperature. A minimum of 500 cells were analyzed across three biological replicates (*n* = 3). Error bars represent the SEM. Log transformation was used to normally distribute the data for statistical analysis (two-way ANOVA; *, *P* < 0.05; **, *P* < 0.01; ns, not significant). 63x scale bar, 10 μm. B. Growth of the WT strain, the *mar1*Δ mutant strain, and the *mar1*Δ + *MAR1* complemented strain was assessed in YPD medium at 37°C. Growth was tracked for 40 hours and was measured by absorbance at OD_600_. Figure summarizes data across three biological replicates (*n* = 3). Error bars represent the SEM. C. Hypoxia resistance was assessed by growth on YES medium in the presence of CoCl_2_ (0.7 mM) and in a microaerophilic chamber. Serial dilutions of the WT strain, the *mar1*Δ mutant strain, the *mar1*Δ + *MAR1* complemented strain, and the *sre1*Δ mutant strain were spotted onto agar plates and incubated at 30°C. Results were compared to the same strains grown in ambient air conditions.

Cell cycle regulation is also known to be related to fungal adaptation to hypoxia (42–44). Because *C. neoformans* is an obligate aerobe, WT fungal cells undergo G_2_-arrest in response to hypoxia (45, 46). We assessed the ability of the *mar1*Δ mutant strain to grow in an environment with reduced oxygen availability by observing growth in the presence of CoCl_2_ and in a microaerophilic chamber. In both cases, we observed that the *mar1*Δ mutant strain displays enhanced growth compared to the WT strain and the *mar1*Δ + *MAR1* complemented strain (Figure 6C). In these assays, the CoCl_2_- and hypoxia-sensitive *sre1*Δ mutant strain was used as a control (31) (Figure 6C). Collectively, these observations suggest that the cell cycle defects of the *mar1*Δ mutant strain may contribute to its ability to survive, slowly proliferate, and persist in the murine granuloma environment.

## DISCUSSION

Here, we report and characterize the host response to a chronic and indolent *C. neoformans* lung infection, one distinguished by sustained granulomas. Using the inhalation route of infection in C57BL/6 mice, we observe granuloma formation in infections due to both the WT and *mar1*Δ mutant strains. However, the appearance, development, and maintenance of these granulomas differ significantly. In WT infections, small, immature granulomas form early in infection. As infection progresses, these nascent granulomas begin to degenerate, leading to fungal proliferation throughout the lungs, fungal dissemination to the brain, and eventually murine death. This type of early, immature granuloma formation has been observed previously in murine infections with other *C. neoformans* WT strains (16, 17). In contrast, in *mar1*Δ mutant strain infections we observe mature pulmonary granulomas that develop over several weeks in the absence of overt clinical symptoms. These granulomas differ from the WT-induced granulomas because they appear later in infection, are typically larger, and are more contained. The containment of these granulomas may be expected because *mar1*Δ-induced granulomas are associated with a significantly lower fungal burden compared to WT strain infections, suggesting that the granulomas effectively inhibit fungal proliferation throughout the lungs. Despite this drastic reduction in fungal burden, the *mar1*Δ mutant strain induces a comparable pulmonary cytokine and leukocyte response to that of the WT strain during early stages of infection. Previous work reported by our group characterized the *mar1*Δ mutant strain as more immunogenic than the WT strain, due to its poorly organized cell wall and impaired polysaccharide capsule attachment (23). We posit that the combination of reduced fungal burden and increased immunogenicity drives *mar1*Δ-induced granuloma formation: the increased immunogenicity results in an immune response that contains the reduced number of *mar1*Δ mutant cells within granulomas during early stages in infection.

We further observe that *mar1*Δ-induced granulomas are maintained throughout infection, from 14 DPI to as late as 100 DPI. We find that the immune microenvironment associated with these granulomas has significantly reduced cytokine and leukocyte responses. Previous work has implicated classically-activated macrophage polarization in enhanced antifungal activity of macrophages (47–49). We find that *mar1*Δ-infected lungs have a comparable number of or fewer (depending on the timepoint) classically-activated (M1) and alternatively-activated (M2) macrophages compared to WT-infected lungs, suggesting that differential polarization of macrophages does not contribute to the reduced fungal burden and associated immune response in *mar1*Δ-infected lungs. Collectively, these observations demonstrate that *mar1*Δ-induced granulomas are largely a fungal-driven phenomenon, with the sustained reduction in *mar1*Δ mutant strain fungal burden resulting in a dampened immune response compared with WT-infected lungs. Using these approaches, we have defined a detailed timeline of granuloma formation, in both WT and *mar1*Δ mutant strain infections, and characterized multiple fungal factors that contribute to granuloma formation (Figure S5).

In addition to the fungal drivers of *mar1*Δ-induced granuloma formation described above, we have also confirmed the role of GM-CSF as a host driver of cryptococcal granuloma formation. From our pulmonary cytokine analyses, we observed that GM-CSF is the only differentially produced cytokine in *mar1*Δ-infected lungs compared to WT-infected lungs. Specifically, GM-CSF is elevated in *mar1*Δ-infected lungs at 3 DPI, an early timepoint in infection at which the pulmonary immune response is being actively developed. This increased GM-CSF production may be a result of increased Dectin-1 activation by the *mar1*Δ mutant strain. We previously reported that the *mar1*Δ mutant strain is partially recognized by the pathogen recognition receptor Dectin-1, likely due its increased exposed surface β-glucan and chitin (23). Dectin-1 has been shown to be required for normal GM-CSF production in murine macrophages (50). Additionally, GM-CSF production is known to result in an increase in Dectin-1 expression by murine macrophages (50, 51). We also report that granuloma formation is dependent on GM-CSF signaling, as granulomas are absent in Csf2rb^-/-^ mouse background infections with either the WT or *mar1*Δ mutant strains. These results are expected because GM-CSF plays a significant role in both *C. gattii* and *C. neoformans* infections, as individuals with GM-CSF autoantibodies are unusually susceptible to cryptococcal infection (52–54). Furthermore, previous work in both mycobacterial (24–26) and cryptococcal infections (17) has demonstrated that GM-CSF signaling is required for granuloma formation, likely due to its requirement for macrophage recruitment to the lung during early stages of infection. Our model enables further exploration of the requirement of GM-CSF for granuloma maintenance. For example, future experiments can introduce GM-CSF antibodies into *mar1*Δ-infected mice to determine whether GM-CSF is required for *mar1*Δ-induced granuloma maintenance and control of infection. Furthermore, WT strain infections can be supplemented with exogenous GM-CSF to determine whether increased GM-CSF can help maintain WT-induced granulomas.

Despite the reduced fungal burden of the *mar1*Δ mutant strain compared to the WT strain, the *mar1*Δ mutant strain persists in the murine lung long-term, up to 100 DPI. Titan cell formation is a well-characterized persistence mechanism that is specific to *Cryptococcus* species (35, 37). Results from an established *in vitro* titanization assay (30) in combination with our histopathological observations demonstrate that the *mar1*Δ mutant strain is unable to form Titan cells, and as a result, Titan cells do not explain the persistence of the *mar1*Δ mutant strain in the murine lung. We also observed that the *mar1*Δ mutant strain is attenuated in the implementation of another important virulence factor, the polysaccharide capsule. Although the *mar1*Δ mutant strain has a similar basal level of capsule to the WT strain, it is unable to extend its capsule to the level of the WT strain in response to capsule-inducing signals.

The expression of many virulence factors is known to be mediated by the cell cycle (41). Furthermore, recent work has proposed that *C. neoformans* undergoes a unique cell cycle *in vivo*, the “stress cell cycle”, that regulates the employment of various virulence factors (55). Titan cells are polyploid cryptococcal cells that form in both human and mouse lungs during infection (35, 37). This polyploidization and concomitant cell body enlargement is negatively regulated by the transcription factor Usv101, which acts downstream of the cell cycle regulator Swi6 (30, 38). Furthermore, recent work has found that the cyclin, Cln1, contributes to Titan cell formation by regulating DNA replication and cell division after G_2_-arrest *in vivo* (55). Similarly, capsule elongation is also regulated by the cell cycle, with the majority of capsule elongation occurring in G_1_ phase of the cell cycle (40). The dysregulation of these cell cycle-mediated virulence factors suggests that the *mar1*Δ mutant strain harbors cell cycle defects.

We indeed observed that the *mar1*Δ mutant strain displays a marked increase in cytokinesis defects compared to the WT strain, at both 30°C and 37°C, leading to a decreased growth rate. In various cell types, including stem cells (56), tumor cells (57), bacteria (58), and fungi (43), a reduction in growth rate is required for survival in the presence of hypoxia. It is possible that its inherent decreased growth rate predisposes the *mar1*Δ mutant strain to growth in a hypoxic environment. The mammalian environment is known to limit oxygen availability to invading microorganisms, as a stressor used to contain microbial proliferation (42). This important resource is likely even further restricted within the pulmonary granuloma, which is known to have suboptimal oxygen levels in the context of mycobacterial infection (59). Recent work by the Alanio laboratory has demonstrated that cryptococcal dormancy can be induced by a combination of nutrient and oxygen deprivation (44, 60). Furthermore, the Dromer laboratory has found that dormant cryptococcal cells are characterized by reduced metabolic activity and delayed growth (43). With these observations in mind, it is possible that the slow growth and hypoxia resistance phenotypes of the *mar1*Δ mutant strain enable its survival and persistence within granulomas in the model described here. Further work will be required to determine whether these phenotypes are necessary and/or sufficient for fungal survival and persistence within granulomas.

The Del Poeta laboratory has developed the most well-characterized murine pulmonary granuloma model of cryptococcal disease to date using the *gcs1*Δ mutant strain. From the fungal perspective, the *gcs1*Δ mutant strain lacks the membrane sphingolipid glucosylceramide, making it an obligate intracellular pathogen and, as a result, completely avirulent in a murine inhalation model, the route of infection that most closely replicates the course of human infection (18, 19). It is noteworthy that both the *gcs1*Δ mutant strain and the *mar1*Δ mutant strain are constructed in the same WT strain background, and as a result, these two mutant strains are comparable and can potentially be used together to explore the complex characteristics of granuloma formation. For example, both strains display cell cycle defects in the presence of physiological stress: the *gcs1*Δ mutant strain arrests at alkaline pH (18) and the *mar1*Δ mutant strain displays cytokinesis defects at 37°C. These similarities suggest that a slow growth phenotype in the host environment may favor fungal containment with granulomas. Virulence potential is a notable difference between the strains. The *gcs1*Δ mutant strain is unable to initiate infection and disease via the inhalation route of infection (18), categorizing *GCS1* as a disease initiation factor (61). In contrast, the *mar1*Δ mutant strain can establish infection and cause fatal disease in nearly half of the infected mice (23), making *MAR1* a disease progression factor (61). This may be related to the fact that *GCS1* orthologs are found in many pathogenic fungi (18), while *MAR1* appears to be a *Cryptococcus*-specific gene (23). These contrasting features suggest that granuloma formation is a highly complex process that relies on the interplay between many fungal and host factors.

From the host perspective, *gcs1*Δ-induced granuloma formation requires host sphingosine kinase 1-sphingosine 1-phosphate (SK1-S1P) signaling (20, 21). Most recently, the Del Poeta laboratory has applied this model to explore cryptococcal reactivation. Mimicking human disease, *gcs1*Δ mutant cells become reactivated from granulomas and disseminate upon immunosuppression with the multiple sclerosis therapeutic FTY720, which suppresses SK1-S1P signaling (22). This model has enabled the first murine reactivation studies of cryptococcal infection. Future work with *mar1*Δ-induced granulomas can similarly explore reactivation in the context of immunosuppression, to better understand the typical course of cryptococcal disease in humans. One of the populations most vulnerable to cryptococcal reactivation includes untreated HIV/AIDS patients (7). In our leukocyte infiltrate analyses, we observed that *mar1*Δ-infected lungs have an enhanced CD4+ T cell response compared to WT-infected lungs at 21 DPI. This observation is particularly striking because *mar1*Δ-infected lungs have a decreased or equivalent response compared to WT-infected lungs for all other leukocytes tested at this same timepoint. CD4+ T cells are present in pulmonary granulomas of immunocompetent individuals (62). Furthermore, CD4+ T cells border the periphery of pulmonary granulomas in HIV+ individuals receiving antiretroviral therapy, but they are lost in individuals with advanced HIV/AIDS, suggesting that CD4+ lymphocytes may be involved in granuloma maintenance (2, 62). By inducing CD4+ T cell depletion, and as a result mimicking the HIV/AIDS disease state, we can probe the role of CD4+ T cells in the maintenance of granulomas in this model. Following immunosuppression, we can observe *mar1*Δ-infected mice to track granuloma breakdown and fungal proliferation with the same approaches used here. Considering both the fungal and host drivers of granuloma formation outlined here, this model harbors features that make it unique from other existing cryptococcal granuloma models.

## ACKNOWLEDGEMENTS

We thank the Duke University School of Medicine for the use of the Research Immunohistochemistry Laboratory Shared Resource, which prepared all histopathology samples. We thank Dr. Joseph Heitman and Anna Floyd-Averette for providing the Csf2rb^-/-^ mice. Flow cytometry was performed in the Flow Cytometry and Laser Capture Microdissection Core Facility at The University of North Texas Health Science Center (UNTHSC) (which is supported by National Institutes of Health award ISIORR018999-01A1) and the Cell Analysis Core at The University of Texas at San Antonio. Scanning electron microscopy was performed at the Chapel Hill Analytical and Nanofabrication Laboratory, CHANL, a member of the North Carolina Research Triangle Nanotechnology Network, RTNN, which is supported by the National Science Foundation, Grant ECCS-1542015, as part of the National Nanotechnology Coordinated Infrastructure, NNCI. This work was supported by R01 AI074677 from the National Institutes of Health to JAA and FLW.

## FIGURE LEGENDS

**Figure S1. Additional histopathology granuloma images.** A. Medium power image from a WT-infected C57BL/6 mouse at 7 DPI, demonstrating a moderate peribronchiolar neutrophilic and mononuclear inflammatory reaction with vague, early, and poorly formed granulomata formation (10X). B. Low power image from a *mar1*Δ-infected C57BL/6 mouse at 3 DPI, demonstrating an absence of a significant inflammatory reaction (4X). C, D. Low power image (C) from a *mar1*Δ-infected C57BL/6 mouse at 40 DPI showing a relatively well-circumscribed nodule containing well developed organizing lymphohistiocytic inflammation and medium power view (D) highlighting compact histiocytic aggregates and peripheral mononuclear cells, characteristic of granuloma formation (C; 4X, D; 10X). E. Medium power image from a WT-infected Csf2rb^-/-^ mouse at 7 DPI showing a marked peribronchiolar neutrophilic and mononuclear inflammatory reaction without granuloma formation (10X). F. Low power image from a WT-infected Csf2rb^-/-^ mouse at 14 DPI demonstrating an absence of a significant inflammatory reaction (4X). All images are of hematoxylin- and eosin-stained tissue sections.

**Figure S2. Complete pulmonary cytokine profile throughout infection.** Pulmonary cytokine responses of female C57BL/6 mice inoculated with 1 × 10^4^ cells of the WT strain or the *mar1*Δ mutant strain were measured using the Bio-Plex protein array system throughout infection: 1 (*n* = 15), 3 (*n* = 15), 7 (*n* = 10), 14 (*n* = 10), and 21 (*n* = 10) DPI. Error bars represent SEM. Statistical significance between strains at each timepoint was determined using Student’s *t* test (*, *P* < 0.05; **, *P* < 0.01; ***, *P* < 0.001; ****, *P* < 0.0001; no designation between strains, not significant).

**Figure S3. Complete pulmonary leukocyte infiltrate response throughout infection.** Pulmonary immune cell infiltrates of female C57BL/6 mice inoculated with 1 × 10^4^ cells of the WT strain or the *mar1*Δ mutant strain were measured by flow cytometry throughout infection: 1, 3, 7, 14, and 21 DPI. Data shown are the mean ± of absolute cell numbers from three independent experiments (*n* = 3) performed using five mice per group per timepoint per experiment. Error bars represent the SEM. Statistical significance between strains at each timepoint was determined using Student’s *t* test (*, *P* < 0.05; **, *P* < 0.01; no designation between strains, not significant).

**Figure S4. Recovery of *mar1*Δ mutant cells from murine lungs at extended timepoints in infection.** Lungs from female C57BL/6 mice infected with the *mar1*Δ mutant strain were harvested at 61 and 100 DPI. Single fungal colonies were isolated on YPD agar plates and subsequently incubated in various conditions that allowed for identification of *mar1*Δ mutant isolates: YPD medium at 30°C, YPD medium at 37°C, YPD medium + nourseothricin (NAT), and YPD medium pH 8.15. The original WT strain (A1) and *mar1*Δ mutant strain (A2) are included as controls in each condition.

**Figure S5. Granuloma formation and maintenance timeline.** Chronological summary of important observations about granuloma formation and maintenance in the WT strain (top) and *mar1*Δ mutant strain (bottom) backgrounds. Cartoons adapted from BioRender.com (2021).

